# GCN5L1 promotes diastolic dysfunction by inhibiting cardiac pyruvate oxidation

**DOI:** 10.1101/2021.09.07.459319

**Authors:** Dharendra Thapa, Paramesha Bugga, Bellina A.S. Mushala, Janet R. Manning, Michael W. Stoner, Brenda McMahon, Xuemei Zeng, Pamela S. Cantrell, Nathan Yates, Bingxian Xie, Lia R. Edmunds, Michael J. Jurczak, Iain Scott

## Abstract

Left ventricular diastolic dysfunction is a structural and functional condition that precedes the development of heart failure with preserved ejection fraction (HFpEF). The etiology of diastolic dysfunction includes alterations in fuel substrate metabolism that negatively impact cardiac bioenergetics, and may precipitate the eventual transition to heart failure. To date, the molecular mechanisms that regulate early changes in fuel metabolism leading to diastolic dysfunction remain unclear. In this report, we use a diet-induced obesity model and quantitative acetylproteomics in aged mice to show that inhibitory lysine acetylation of the pyruvate dehydrogenase (PDH) complex promotes energetic deficits and diastolic dysfunction in mouse hearts. Cardiomyocyte-specific deletion of the mitochondrial lysine acetylation regulatory protein GCN5L1 prevented hyperacetylation of the PDH complex subunit PDHA1, allowing aged obese mice to continue using pyruvate as a bioenergetic substrate in the heart. Our findings suggest that changes in mitochondrial protein lysine acetylation represent a key metabolic component of diastolic dysfunction that precedes the development of heart failure.

## INTRODUCTION

Left ventricular diastolic dysfunction is an independent pre-clinical predictor of all-cause mortality ^[1]^, and is a prerequisite for the diagnosis of heart failure with preserved ejection fraction (HFpEF) ^[2]^. While the etiology of the diastolic dysfunction is multifactorial, deleterious changes in cardiac energy metabolism underpin the development of both diastolic dysfunction and HFpEF ^[3,4]^. Studies on two recently developed murine models of HFpEF have reported maladaptive changes to the relative oxidation rates of fatty acids, ketones, and glucose in failing hearts ^[5,6]^, suggesting that this metabolic phenotype is central to the development of cardiac dysfunction. Interestingly, the HFpEF phenotype in these two independent mouse models is also characterized by an increase in mitochondrial protein acetylation, a reversible posttranslational modification that is often linked to altered metabolic enzyme activity ^[5,6]^. Restoring normal mitochondrial protein acetylation through supplementation with either nicotinamide riboside (to stimulate SIRT3-mediated protein deacetylation) ^[5]^, or ketone bodies (which inhibit fatty acid uptake and utilization) ^[6]^, resulted in normalized cardiac bioenergetics and amelioration of the HFpEF phenotype ^[5,6]^.

These key studies suggest that tight regulation of mitochondrial protein acetylation is required to prevent the development of HFpEF, and therefore approaches that reduce maladaptive protein hyperacetylation are potentially cardioprotective. We have previously shown that diet-induced increases in the abundance of a mitochondrial lysine acetylation regulatory protein, GCN5L1, correlated with increased fatty acid oxidation and reduced glucose utilization in the heart ^[7,8]^. Here, using a mouse model of constitutively reduced mitochondrial protein acetylation (cardiomyocyte-specific deletion of GCN5L1) and quantitative acetylproteomics, we examined the metabolic mechanisms underlying diastolic dysfunction in mice prior to the development of heart failure. Our results suggest that long-term exposure to nutrient excess in aged mice promotes diastolic dysfunction in the heart, via inhibitory acetylation of enzymes involved in cardiac pyruvate oxidation.

## METHODS

### Transgenic Mice

C57BL/6NJ wildtype (GCN5L1^WT/WT^,Cre^+/-^; “WT”) and cardiomyocyte-specific inducible GCN5L1 knockout (GCN5L1^FL/FL^, Cre^+/-^; “KO”) mice used in the studies were generated as previously reported ^[9]^. GCN5L1 deletion in cardiomyocytes was induced via single tamoxifen injection (40 mg/kg IP) as previously described ^[9]^.

### Animal Care and Experimental Diets

Male WT and GCN5L1 KO animals aged 5-7 months were fed either a standard low fat diet (LFD; Research Diets D12450B), or a high fat diet (HFD; Research Diets D12492), for 30 weeks. Mice were euthanized by CO_2_ asphyxiation and rapid cervical dislocation. All animal procedures were approved by the University of Pittsburgh Institutional Animal Care and Use Committee.

### Quantitative Acetylproteomics

Acetylated peptides from flash-frozen left ventricle were isolated the acetyl PTMScan Kit (Cell Signaling). Peptides were subject to LC-MS/MS, and the data analyzed using the MaxQuant software suite and Matlab-based script developed in-house.

### Respirometry

Respirometry was performed using an Oroboros O2K High-Resolution Respirometer using mitochondrially-enriched cardiac samples. Protein concentration of the mitochondrially-enriched fraction was determined by BCA protein assay, and oxygen consumption or flux expressed per mg protein normalized to citrate synthase activity.

### Cell Culture and Transfection

Stable control shRNA and GCN5L1 shRNA AC16 cells were described previously ^[10]^. WT PDHA1-Myc, 5KR PDHA1-Myc, and 5KQ PDHA1-Myc were produced using custom gene synthesis (Genscript), and cloned into the pCMV-3Tag-4a plasmid vector. Cells were transfected for 24 h using X-tremeGENE reagent.

### Protein Isolation, Western Blotting, and Immunoprecipitation

For western blotting, tissues were minced and lysed in CHAPS buffer (1% CHAPS, 150 mM NaCl, 10 mM HEPES, pH 7.4). Protein expression was analyzed using the following primary antibodies; rabbit PDHA1 Ser-293 antibody (Cell Signaling; #31866), rabbit PDHA1 (Cell Signaling; #3205), and rabbit GAPDH (Cell Signaling; #2118). GCN5L1 antibody was made by Covance and previously validated ^[11]^. For immunoprecipitation experiments, tissues were minced and lysed in CHAPS buffer. Protein lysates were incubated overnight at 4 °C with rabbit acetyl-lysine (Ac-K, #9441) from Cell Signaling Technology. Protein densitometry was measured using Image J software (National Institutes of Health, Bethesda, MD).

### Statistics

Graphpad Prism software was used to perform statistical analyses. Means ± SEM were calculated for all data sets. Data were analyzed using two-way ANOVA with Tukey’s post-hoc multiple comparison testing to determine differences between genotypes and feeding groups. Data were analyzed with two-tailed Student’s T-tests to determine differences between single variable groups. *P* < 0.05 was considered statistically significant.

## RESULTS

### Cardiomyocyte-specific GCN5L1 cKO mice are resistant to diastolic dysfunction induced by a high fat diet

While the two HFpEF mouse studies referenced above used diet-based two-hit ^[5]^ or three-hit ^[6]^ models to produce the HFpEF phenotype, long-term exposure to a high fat diet (HFD) alone can promote diastolic dysfunction ^[12]^. We therefore used a long-term HFD model in aged mice to specifically examine diastolic dysfunction in a pre-HFpEF state. Wildtype (WT) and cardiomyocyte-specific GCN5L1 knockout (KO) mice (which display global reductions in mitochondrial protein acetylation) ^[9]^ were placed on low fat (LFD; 10% fat) or high fat (HFD; 60% fat) diets for 30 weeks (**Figure 1A**), and heart structure and function were measured by ultrasonography (**Figure 1B, Figure S1**). No significant changes in left ventricle mass, cardiac output, or systolic function were observed between WT and KO mice on a HFD (**Figure 1C-E**). In contrast, WT mice displayed a significant increase in E/e’ ratio (a marker of diastolic dysfunction) ^[5]^ on a HFD relative to acetylation-deficient KO mice under the same conditions (**Figure 1F**). Combined, these data suggest that GCN5L1 expression in cardiomyocytes has a specific negative effect on cardiac diastolic function in aged mice after exposure to a long-term HFD.

**Figure 1:**
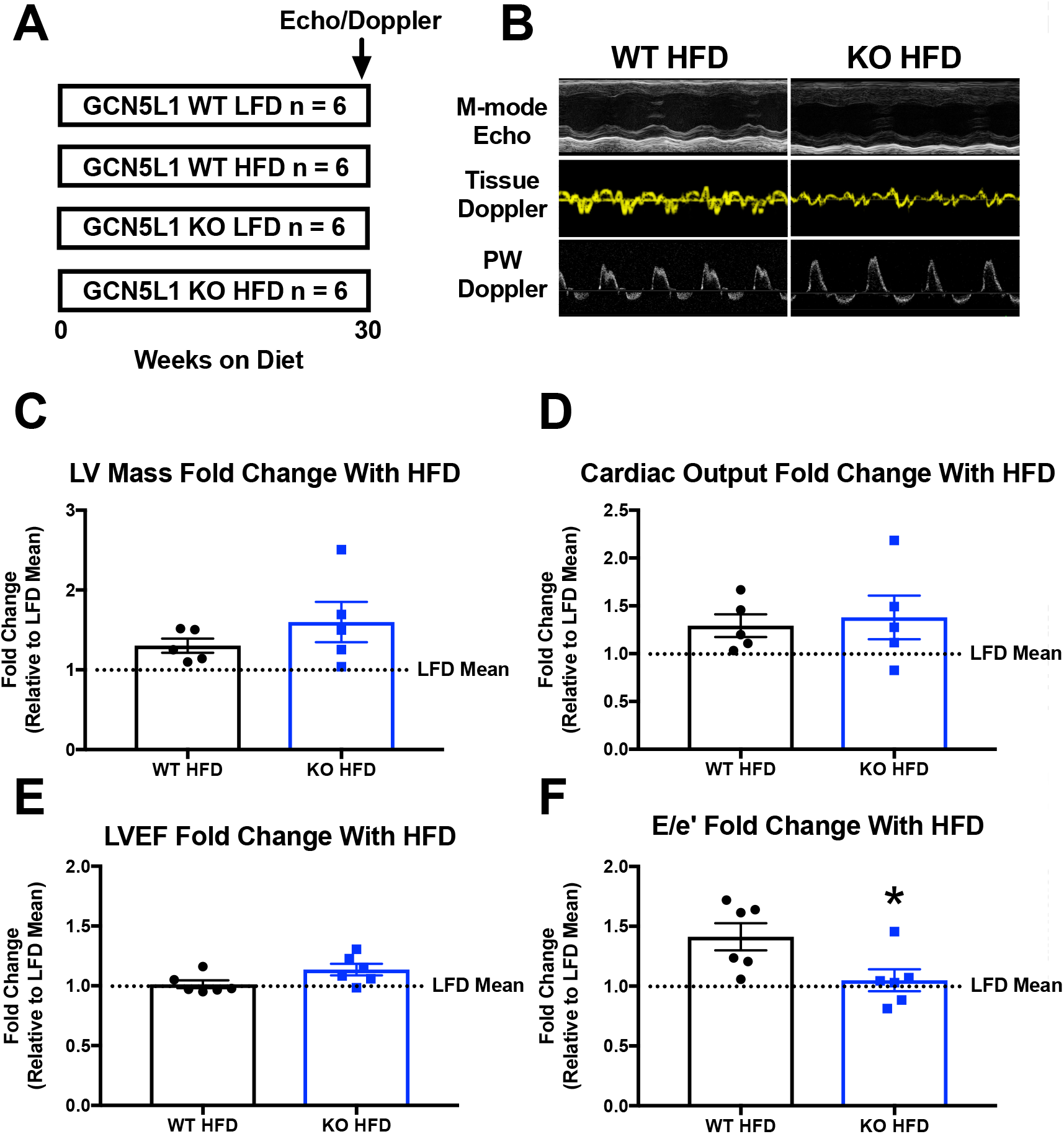
GCN5L1 promotes diastolic dysfunction in aged mice after exposure to a long-term high fat diet. **A.** Wildtype (WT) and cardiomyocyte-specific GCN5L1 knockout (KO) mice aged 5-7 months were placed on low fat (LFD), or high fat diets (HFD) for 30 weeks. **B-F.** Ultrasonography measurements of left ventricle (LV) mass, cardiac output, systolic (left ventricle ejection fraction; LVEF), and diastolic function (mitral valve E/e’ wave ratio). Data are presented as fold change in HFD mice relative to LFD genotype controls. Full ultrasonography data are shown in Table S1. N = 6; T-test; * = *P* < 0.05.

### GCN5L1 cKO mice have enhanced cardiac pyruvate oxidation capacity under high fat diet conditions

To understand which mitochondrial metabolic pathways are regulated by GCN5L1-mediated acetylation under conditions that promote diastolic dysfunction, we performed quantitative acetylproteomics on HFD-fed WT and KO mice (**Figure 2A**). Gene ontology analysis using PANTHER demonstrated that enzymes involved in mitochondrial fatty acid oxidation and pyruvate metabolism were among the top three pathways affected by increased GCN5L1-mediated lysine acetylation (**Figure 2B; Table S2**).

**Figure 2:**
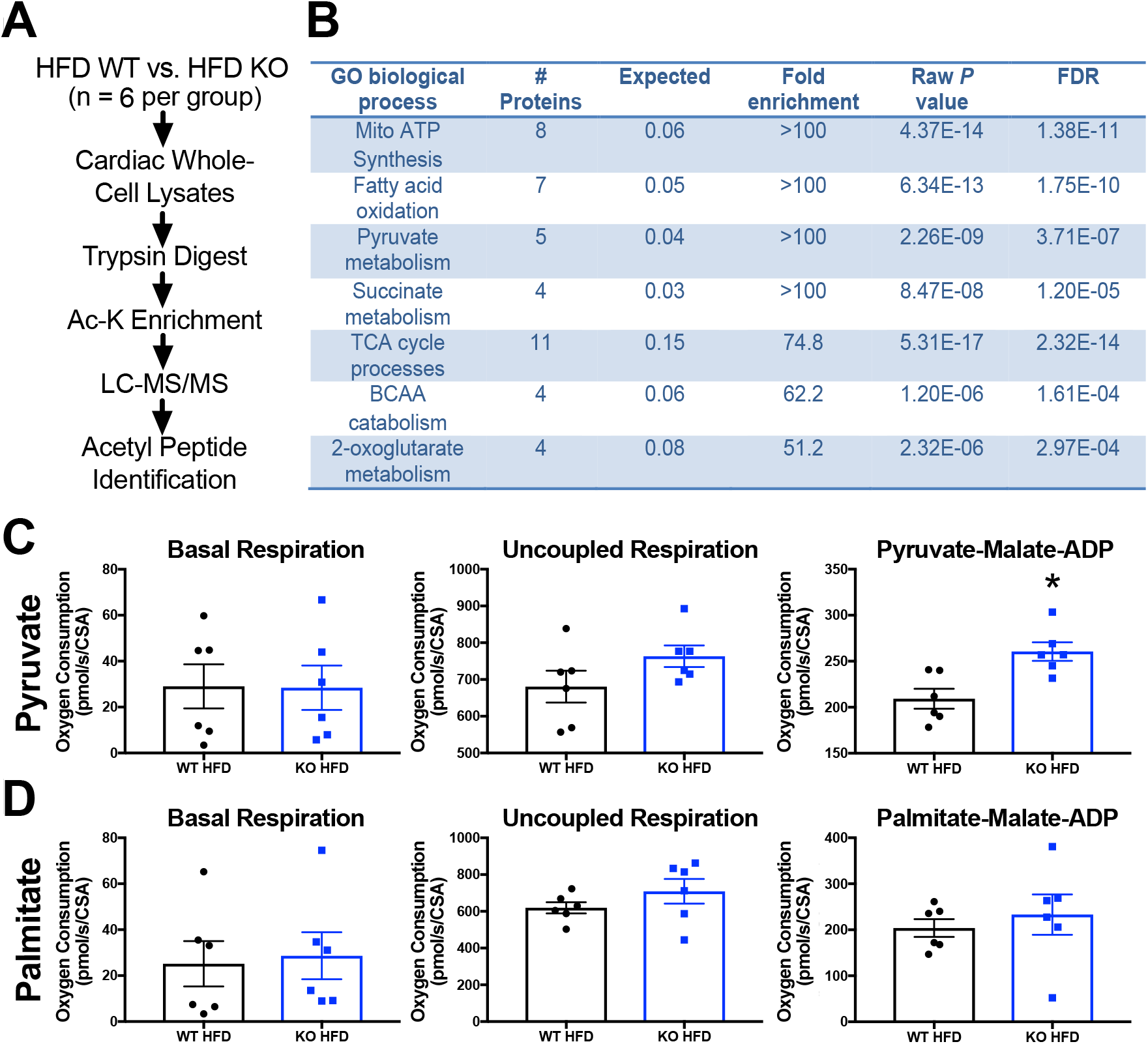
GCN5L1 inhibits cardiac pyruvate oxidation in obese mice. **A-B.** Quantitative acetylproteomics were used to identify biological pathways that were subject to GCN5L1-mediated lysine acetylation in high fat diet (HFD) mouse hearts. GO enrichment analysis, *P* values, and False Discovery Rates (FDR) were calculated using PANTHER. A full list of proteins from each pathway is shown in Table S2. **C-D.** Oxygen consumption in cardiac tissues from HFD-fed WT and KO mice was measured in response to either pyruvate-malate-ADP (**C**) or palmitate-malate-ADP (**D**) using Oroboros respirometry. N = 6; T-test; * = *P* < 0.05.

To understand if one or both fuel metabolism pathways were disrupted in mice with diastolic dysfunction, we performed Oroboros respirometry in cardiac tissue lysates from HFD-fed WT and KO mice. Previous reports have shown that fatty acid oxidation is significantly modified in hyperacetylated mitochondria from HFpEF-model mice ^[5,6]^. However, we found here that ADP-stimulated state 3 respiration was not significantly different between HFD WT and KO samples provided with palmitate as the fuel substrate. Instead, we observed a significant increase in ADP-stimulated state 3 respiration in HFD KO tissues given pyruvate as a fuel source, relative to their WT counterparts on the same diet (**Figure 2C-D; Figure S1**). Combined, these data indicate that a loss of pyruvate oxidation, rather than aberrant fatty acid oxidation, underpins the energetic deficits that contribute to the development of diastolic dysfunction in this model.

### Exposure to a high fat diet inhibits pyruvate oxidation through hyperacetylation of PDHA1

The pyruvate dehydrogenase (PDH) complex regulates mitochondrial pyruvate utilization, and previous work has demonstrated that increasing PDH activity can reverse diastolic dysfunction in diabetic rodents ^[13,14]^. Elevated PDH activity is commonly associated with a decrease in inhibitory phosphorylation at Ser-293 ^[5]^, however we detected no difference in PDH phosphorylation between WT and KO mice on either diet (**Figure 3A**). Instead, we found a significant increase in the acetylation status of PDHA1 (a component of the PDH E1 enzyme) in HFD-fed WT mice relative to LFD controls, and a significant decrease in PDHA1 acetylation in both KO diet groups (**Figure 3B**). Consistent with this, PDH activity was significantly increased in KO mice on a long-term HFD relative to WT mice (**Figure 3C**), and PDHA1 acetylation status across all animal groups was negatively correlated with PDH activity by linear regression (R^2^ = 0.3678; *P* = 0.037; **Figure 3D**). Combined, these data suggest that exposure to a long-term HFD in aged mice inhibits PDH through acetylation, rather than the more common phosphorylation inhibitory pathway.

**Figure 3:**
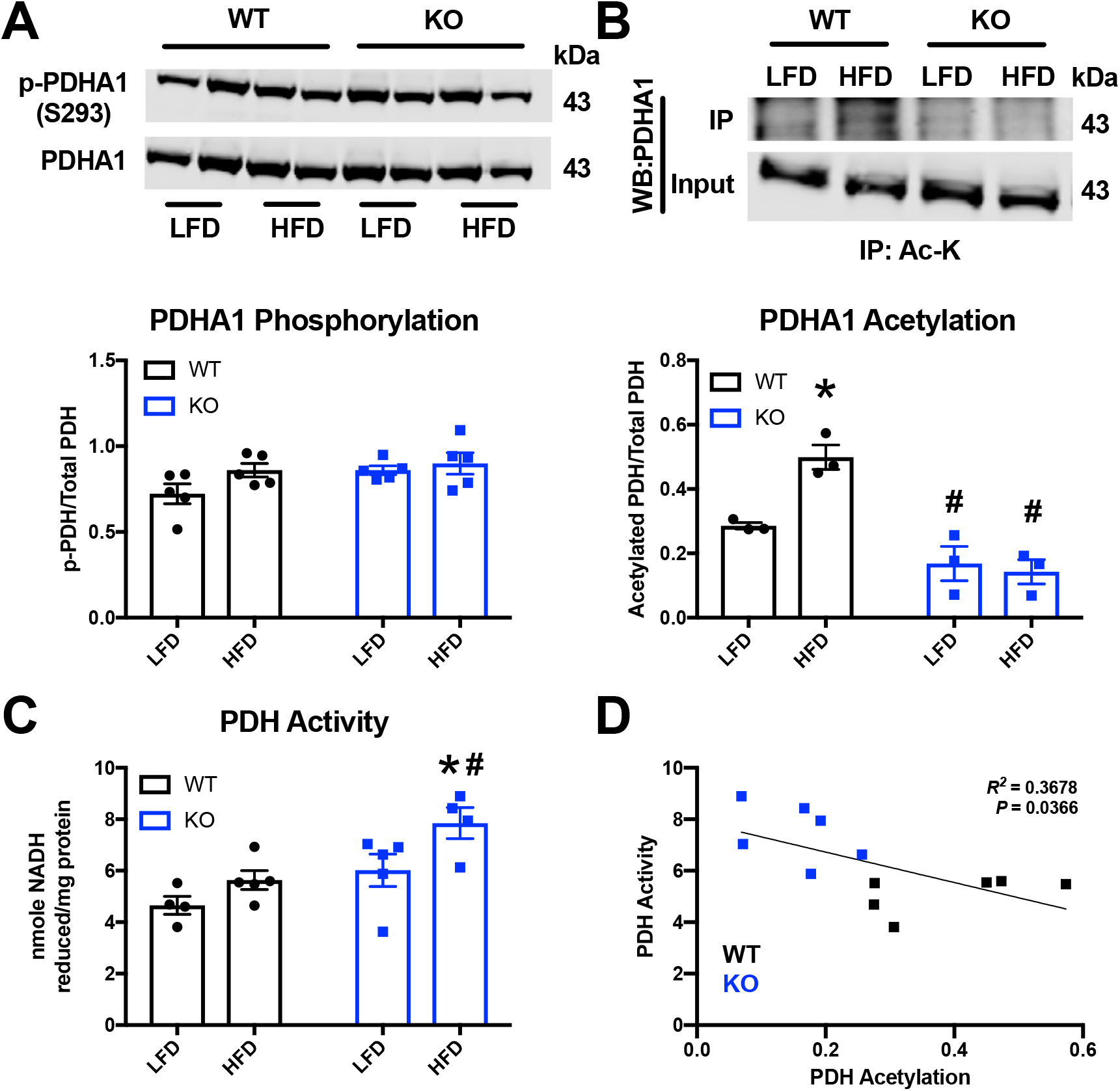
GCN5L1 promotes PDHA1 acetylation and inhibits pyruvate dehydrogenase activity. **A-B.** PDHA1 phosphorylation (S293) and lysine acetylation in WT and KO mice following LFD or HFD. N = 3-5; Two-way ANOVA; * = *P* < 0.05 relative to WT LFD, # = *P* < 0.05 to WT HFD **C.** PDH activity in WT and KO mice following LFD or HFD. N = 3-5; Two-way ANOVA; * = *P* < 0.05 relative to WT LFD, # = *P* < 0.05 to WT HFD. **D.** Linear regression of PDH activity and acetylation levels in WT and KO mice following LFD or HFD.

### Preventing PDHA1 acetylation promotes pyruvate dehydrogenase activity in cardiac cells

To verify these findings independently, we examined PDH activity and acetylation status in the human-derived AC16 cardiomyocyte cell model following shRNA-mediated GCN5L1 depletion. Knockdown of GCN5L1 in AC16 cells resulted in a ~40% increase in PDH activity, and a ~50% decrease in PDHA1 acetylation, without observed changes in total PDH abundance (**Figure 4A-C**). Combined with our *in vivo* findings, these data suggest that GCN5L1-mediated hyperacetylation of PDH is a key driver of defects in pyruvate oxidation in cardiac cells.

**Figure 4:**
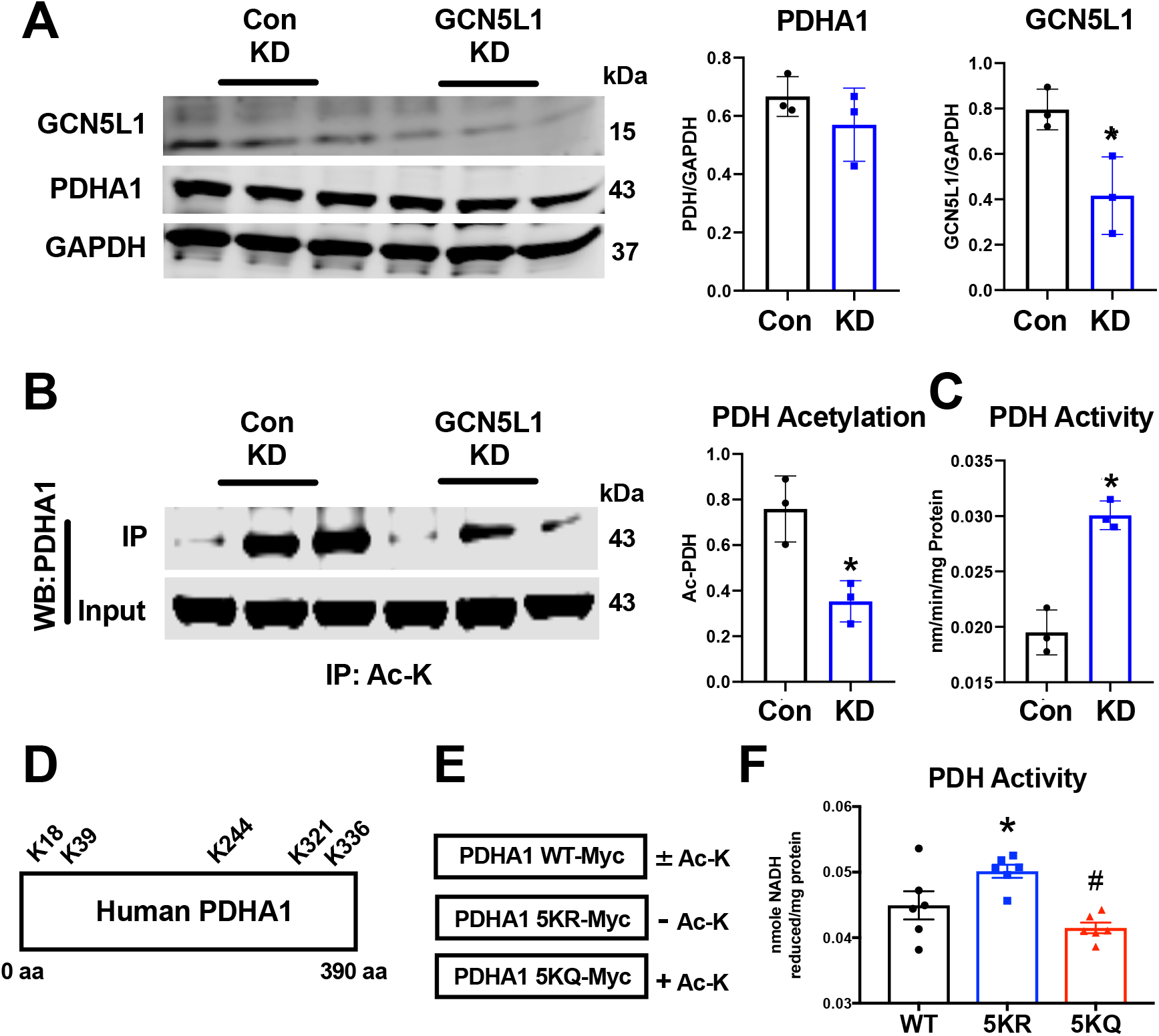
PDHA1 lysine acetylation directly inhibits PDH activity in cardiac cells. **A-C.** Total PDH acetylation and enzymatic activity was measured in stable Control shRNA or GCN5L1 shRNA knockdown (KD) human cardiac AC16 cells. N = 3; T-test; * = *P* < 0.05 relative to WT. **D.** Schematic of human PDHA1 showing published lysine acetylation sites. **E-F.** GCN5L1 KD AC16 cells were transfected with Myc-tagged wildtype PDHA1 (WT-Myc), a non-acetylated PDHA1 mimic in which lysines K18/K39/K244/K321/K336 were replaced with arginine (5KR-Myc), and an acetylated PDHA1 mimic where lysines K18/K39/K244/K321/K336 were replaced with glutamine (5KQ-Myc). N = 6; One-way ANOVA; * = *P* < 0.05 relative to WT, # = *P* < 0.05 to 5KR.

Finally, we sought to determine the specific effect of lysine hyperacetylation on PDH enzymatic activity. Published studies in the PhosphoSitePlus proteomics database (www.phosphosite.org) have identified five acetylation sites on human PDHA1 at K18, K39, K244, K321, and K336 (**Figure 4D**). We generated novel PDHA1 expression constructs, where each of these five lysines were replaced with either arginine (5KR; to mimic a non-acetylated state) or glutamine (5KQ; to mimic a hyperacetylated state; **Figure 4E**), and expressed them in GCN5L1 knockdown AC16 cells (which represent a basal, non-acetylated state for endogenous PDHA1). PDH activity was significantly increased in non-acetylated PDHA1 5KR expressing cells relative to those overexpressing wildtype PDHA1 (**Figure 4F**). Conversely, PDH activity was significantly decreased in hyperacetylated PDHA1 5KQ expressing cells relative to those deficient in PDHA1 acetylation (**Figure 4F**). Together, these *in vitro* data indicate that hyperacetylation of PDHA1 has a specific negative effect on PDH enzymatic activity in cardiac cells.

## DISCUSSION

In conclusion, we show that inhibitory acetylation of PDHA1 drives reduced pyruvate utilization in a diet-based model of cardiac diastolic dysfunction. Mice lacking cardiomyocyte-expression of the mitochondrial acetylation regulator GCN5L1 are protected from diastolic dysfunction, which is linked to an absence of inhibitory PDHA1 acetylation.

Previous work from our group and others has demonstrated that increased mitochondrial protein acetylation in the heart is linked with increased rates of fatty acid oxidation *ex vivo* and *in vitro* ^[7,15,16]^. However, the functional consequence of this change is unclear, as increased fatty acid oxidation activities from hyperacetylated hearts did not lead to increased contractility when measured *ex vivo* ^[17]^. Results from the current study are instead in agreement with previous work showing that PDH acetylation is linked to a decrease in its activity in the heart *in vivo* and *ex vivo* ^[8,18]^, and that GCN5L1 overexpression leads to elevated PDH acetylation and *in vitro* ^[8]^. This is the first loss-of-function study showing that GCN5L1 regulates the acetylation status of PDHA1 *in vivo*, and demonstrates that increased abundance of this modification is linked to both decreased pyruvate utilization, and impaired diastolic function, in the hearts of aged, obese mice. Our work suggests that manipulation of PDHA1 acetylation levels *in vivo* may represent a novel target for therapeutic intervention in the treatment of diastolic dysfunction.

The inhibitory effect of PDH acetylation in diastolic dysfunction described here is consistent with its role in mouse models of heart failure with reduced ejection fraction (HFrEF), where miR-195-mediated downregulation of the deacetylase SIRT3 leads to increased acetylation of PDH subunits and decreased PDH enzymatic activity ^[19]^. However, in this study we detected no differences in SIRT3 abundance in response to diet or genotype (**Figure S3**), placing GCN5L1 at the center of mitochondrial acetylome regulation in this diastolic dysfunction model.

Interestingly, the deletrious effect of global mitochondrial protein hyperacetylation in the development of HFpEF ^[5,6]^ sits at odds with its demonstrated lack of effect in mouse surgical models of HFrEF ^[20,21]^. In this study, mice with a combined genetic deletion of the deacetylase, SIRT3, and the carnitine acetyltransferase, CrAT, did not show increased susceptibility to pressure overload-induced heart failure despite massively hyperacetylated mitochondria ^[20]^. Further work will be required to determine whether the differences found reflect the etiology of the different models used (i.e. nutrition vs. surgical pressure overload); the site-specific nature of the acetylation sites modified on each protein (i.e. whether lysine residues on the same protein are differentially regulated by the two models); or whether the different mechanisms by which GCN5L1 and SIRT3 control mitochondrial protein acetylation status (acetyl-CoA dependent acetylation vs. NAD^+^ dependent deacetylation), lead to different functional outcomes.

## SOURCES OF FUNDING

This work was supported by: NIH/NHLBI K99/R00 (HL146905) to DT; NIGMS T32 (GM133332) to B.A.S.M; and NIH/NHLBI R01 (HL132917 & HL147861) research grants to IS. The University of Pittsburgh Center for Metabolism and Mitochondrial Medicine is supported by a Pittsburgh Foundation (MR2020 109502) grant to MJJ. The University of Pittsburgh Rodent Ultrasonography Core received support from the NIH/OD S10 Instrumentation Program (OD023684).

## DISCLOSURES

None

## AUTHOR CONTRIBUTIONS

**Performed experiments:** DT, PB, BASM, JRM, MWS, BM, XZ, PC, BX, LRE. **Designed experiments:** DT, PB, NY, MJJ, IS. **Analyzed data:** DT, PB, BM, XZ, PC, NY, BX, LRE, MJJ, IS. **Produced figures:** DT, PB, IS. **Wrote/edited manuscript:** DT, PB, NY, MJJ, IS.

## Notes

### Competing Interest Statement

The authors have declared no competing interest.

### Summary of Updates

We have updated the figures, added new data, and provided more information on experimental methods.

