## Supplemental Information for "GCN5L1 promotes diastolic dysfunction by inhibiting cardiac pyruvate oxidation"

**Supplemental Methods**

**Animal Care and Experimental Diets**

Animals were housed in the University of Pittsburgh animal facility under standard conditions with *ad libitum* access to water and food, and maintained on a constant 12h light/12h dark cycle. Male WT and GCN5L1 KO animals aged 5-7 months were fed either a standard low fat diet (LFD; Research Diets D12450B), or a high fat diet (HFD; Research Diets D12492), for 30 weeks. Mice were euthanized by $CO_2$ asphyxiation and rapid cervical dislocation. All animal procedures were approved by the University of Pittsburgh Institutional Animal Care and Use Committee.

**Protein Isolation, Western Blotting, and Immunoprecipitation**

For western blotting, tissues were minced and lysed in CHAPS buffer (1% CHAPS, 150 mM NaCl, 10 mM HEPES, pH 7.4) on ice for ~2 hours. Homogenates were spun at 10,000 *g*, and supernatants were collected. Protein lysates were prepared in LDS sample buffer, separated using Bolt SDS/PAGE 4-12% or 12% Bis-Tris gels, and transferred to nitrocellulose membranes (all Life Technologies). Protein expression was analyzed using the following primary antibodies; rabbit PDHA1 Ser-293 antibody (Cell Signaling; #31866), rabbit PDHA1 (Cell Signaling; #3205), and rabbit GAPDH (Cell Signaling; #2118). GCN5L1 antibody was made by Covance and previously validated [9]. Fluorescent anti-mouse or anti-rabbit secondary antibodies (red, 700 nm; green, 800 nm) from Li-Cor were used to detect expression levels. Protein densitometry was measured using Image J software (National Institutes of Health, Bethesda, MD). For immunoprecipitation experiments, tissues were minced and lysed in CHAPS buffer (1% CHAPS, 150 mM NaCl, 10 mM HEPES, pH 7.4) on ice for ~2 hours. Homogenates were spun at 10,000 *g*, and supernatants were collected. Protein lysates were incubated overnight at 4 ºC with rabbit acetyl-lysine (Ac-K, #9441) from Cell Signaling Technology. Immunocaptured proteins were isolated using Protein-G agarose beads (Cell Signaling Technology, #9007), washed multiple times with CHAPS buffer, and then eluted in LDS sample buffer (Life Technologies) at 95 ºC. Samples were separated on 12% Bis-Tris Bolt gels and probed with appropriate antibodies. Protein densitometry was measured using Image J software (National Institutes of Health, Bethesda, MD).

**Ultrasonography**

Animals were anesthetized using isofluorane (1.5-2.0 % v/v by inhalation) and monitored for cardiac functional parameters using a VisualSonics Vevo 3100 in M-mode echocardiography, Tissue Doppler, and Pulsed-Wave Doppler modes. Markers of systolic and diastolic function were calculated using standard echocardiography equations.

**Respirometry**

Citrate synthase activity was determined by preparing a mitochondrially-enriched supernatant by homogenizing a 50 mg piece of left ventricle in 1 ml of STE buffer (250 mM sucrose, 10 mM Tris-HCl, 1 mM EDTA) with a glass vessel and Teflon pestle, followed by centrifugation at 800 *g* for 10 min at 4 °C. Citrate synthase activity was then measured by mixing the mitochondrially-enriched fraction with citrate synthase activity buffer (0.25% Triton X-100, 0.31 mM acetyl-CoA, 0.1 mM 5,5'-dithiobis(2-nitrobenzoic acid), 0.1 M triethanolamine, 1 mM EDTA, and 1 M Tris-HCl), followed by addition of 5 mM oxaloacetate to initiate the reaction. The change in absorbance at 412 nm was recorded every 10 seconds over a two-min period using a spectrophotometer set to 37 $^{o}$C, and activity was calculated as the slope from the linear portion of the curve and normalized to the protein concentration determined by BCA protein assay. Citrate synthase activity was used to normalize respirometry measurements. Respirometry was performed using an Oroboros O2K High-Resolution Respirometer using mitochondrially-enriched cardiac samples prepared as described above. Respiration was measured in MiR05 buffer at 37 $^{o}$C under constant mixing in a sealed, 2 ml chamber. The respirometry protocol consisted of sequential additions of substrates and inhibitors as follows: pyruvate (5 mM) or BSA-conjugated palmitate (0.25 mM), malate (2 mM), ADP (2 mM); oligomycin (2.5 μM); carbonyl cyanide 4-(trifluoromethoxy)phenylhydrazone (FCCP; titrations of 0.5 μM until maximal respiration reached); antimycin A (2.5 μM). Protein concentration of the mitochondrially-enriched fraction was determined by BCA protein assay, and oxygen consumption or flux expressed per mg protein normalized to citrate synthase activity.

**Quantitative Acetylproteomics**

Flash-frozen left ventricle tissues were homogenized in sodium deoxycholate (SDC) lysis buffer using a bead mill (3 x 60 s at 6.5 s/m), and sonicated for 15 minutes in a water bath to shear DNA. Extracted proteins (1.5 mg) were reduced (DTT) and alkylated (CAA), the SDC buffer removed by acid precipitation, and desalted with Sep-Pek C18. After in-solution trypsin digest, 95% of the sample was enriched using the acetyl PTMScan Kit (Cell Signaling). Peptides were eluted twice, and desalted using PepClean C18 (Pierce). Peptides were subject to LC/MS/MS, and the data analyzed using the MaxQuant software suite and Matlab-based script developed in-house. The ratio of acetyl-peptide enrichment between WT HFD and KO HFD samples was determined for each detected mitochondrial peptide, and all peptides where WT acetylation was greater than KO acetylation were selected (Table S1). Identified genes corresponding to these peptides were entered into the PANTHER gene ontology search engine (www.geneontology.org), and grouped according to functional class. Functional classes containing ≥ 4 independent genes were selected for analysis. *P* values, fold enrichment, and false discovery rates were determined by PANTHER.

**Biochemical Assays**

PDH activity was assessed using a commercial kit (MAK-183; Sigma-Aldrich) according to the manufacturer's instructions.

### Supplemental Data

| Parameter | WT LFD | KO LFD | WT HFD | KO HFD |
|---|---|---|---|---|
| Heart Rate (BPM) | 463 ± 51 | 476 ± 40 | 494 ± 50 | 475 ±22 |
| Volume (s; μL) | 28 ± 11 | 20 ± 8 | 31 ± 6 | 38 ± 20 |
| Volume (d; μL) | 67 ± 13 | 62 ± 28 | 79 ± 15 | 93 ± 39 |
| LV Mass (mg) | 140 ± 27 | 159 ± 44 | 179 ± 27 * | 238 ± 80 |
| Cardiac Output (mL/min) | 19 ± 5 | 20 ± 11 | 23 ± 5 | 26 ± 10 |
| LV AW (s; mm) | 1.2 ± 0.2 | 1.5 ± 0.1 | 1.3 ± 0.1 | 1.5 ± 0.1 |
| LV AW (d; mm) | 0.9 ± 0.1 | 1.1 ± 0.1 | 1.1 ± 0.1 * | 1.2 ± 0.1 |
| LV PW (s; mm) | 1.2 ± 0.3 | 1.4 ± 0.2 | 1.4 ± 0.2 | 1.5 ± 0.2 |
| LV PW (d; mm) | 0.9 ± 0.2 | 1.0 ± 0.1 | 1.0 ± 0.1 | 1.1 ± 0.1 |
| Stroke Volume (μL) | 40 ± 8 | 42 ± 21 | 48 ± 11 | 54 ± 20 |
| LV EF (%) | 60 ± 12 | 67 ± 7 | 61 ± 5 | 60 ± 6 |
| LV FS (%) | 31 ± 8 | 37 ± 6 | 32 ± 4 | 32 ± 4 |
| E (mm/s) | 521 ± 65 | 636 ± 41 | 645 ± 87 * | 672 ± 101 |
| e' (mm/s) | 27 ± 4 | 27 ± 5 | 24 ± 4 | 28 ± 6 |
| E/e' (ratio) | 20 ± 3 | 24 ± 5 | 28 ± 5 * | 26 ± 5 |

BPM – Beats Per Minute; S – Systolic; D – Diastolic; LV – Left Ventricle; AW – Anterior Wall; PW – Posterior Wall; EF – Ejection Fraction; FS – Fractional Shortening; E – Mitral Valve Early Filing Velocity; E' – Early Diastolic Mitral Annular Velocity. * = $P < 0.05$ vs. WT LFD

**Table S1 – Ultrasonography parameters in WT and GCN5L1 KO mice after long-term LFD or HFD exposure.** N = 6 per group.

| Biological Process | Acetylated Proteins (WT>KO) |
|---|---|
| Mito ATP Synthesis | Atp5a1, Atp5o, Atp5c1, Atp5j2, mt-Atp8, Atp5j, Atp5f1, Atp5d |
| Fatty acid oxidation | Etfdh, Ivd, Acads, Etfb, Acadvl, Etfa, Acadm |
| Pyruvate metabolism | Pdha1, Dlat, Dld, Pdhx, Pdhb |
| Succinate metabolism | Aldh5a1, Sucla2, Suclg1, Sdha |
| TCA cycle processes | Pdha1, Sucla2, Dlat, Suclg1, Idh3a, Sdha, Idh2, Pdhb, Mrps36, Cs, Mdh2, Aco2 |
| BCAA catabolism | Aldh6a1, Hibch, Mcc2, Ivd |
| 2-oxoglutarate metabolism | Dld, Idh3a, Mrps36, Got2 |

**Table S2 – Mitochondrial proteins with increased lysine acetylation in HFD WT hearts relative to HFD KO hearts.** N = 6 per group.

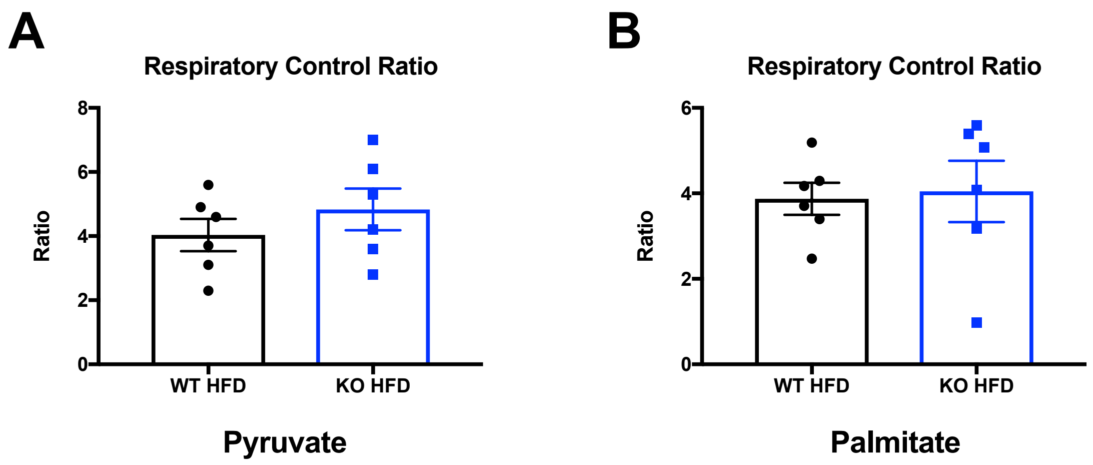


**Figure S1 – Respiratory control ratios of HFD WT and GCN5L1 KO mice provided with either pyruvate or palmitate as a fuel source in Oroboros respirometry studies.** N = 6 per group.

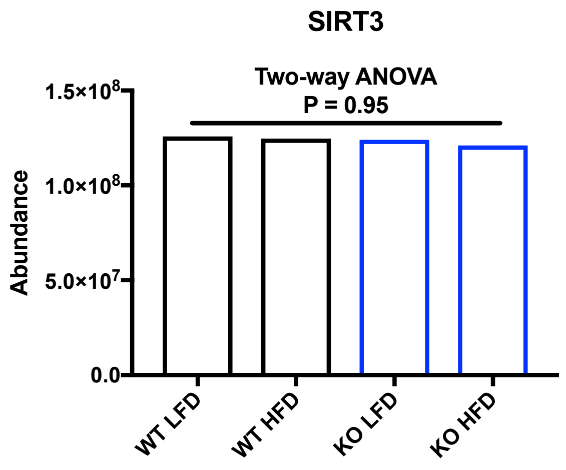


**Figure S2 – SIRT3 protein abundance in WT and GCN5L1 KO mice after long-term LFD or HFD exposure.** N = 6 per group.
